# Interactome dynamics during heat stress signal transmission and reception

**DOI:** 10.1101/2024.04.29.591712

**Authors:** Sung-Gun Park, Andrew Keller, Nathan K. Kaiser, James E. Bruce

## Abstract

Among evolved molecular mechanisms, cellular stress response to altered environmental conditions to promote survival is among the most fundamental. The presence of stress-induced unfolded or misfolded proteins and molecular registration of these events constitute early steps in cellular stress response. However, what stress-induced changes in protein conformations and protein-protein interactions within cells initiate stress response and how these features are recognized by cellular systems are questions that have remained difficult to answer, requiring new approaches. Quantitative *in vivo* chemical cross-linking coupled with mass spectrometry (qXL-MS) is an emerging technology that provides new insight on protein conformations, protein-protein interactions and how the interactome changes during perturbation within cells, organelles, and even tissues. In this work, qXL-MS and quantitative proteome analyses were applied to identify significant time-dependent interactome changes that occur prior to large-scale proteome abundance remodeling within cells subjected to heat stress. Interactome changes were identified within minutes of applied heat stress, including stress-induced changes in chaperone systems as expected due to altered functional demand. However, global analysis of all interactome changes revealed the largest significant enrichment in the gene ontology molecular function term of RNA binding. This group included more than 100 proteins among multiple components of protein synthesis machinery, including mRNA binding, spliceosomes, and ribosomes. These interactome data provide new conformational insight on the complex relationship that exists between transcription, translation and cellular stress response mechanisms. Moreover, stress-dependent interactome changes suggest that in addition to conformational stabilization of RNA-binding proteins, adaptation of RNA as interacting ligands offers an additional fitness benefit resultant from generally lower RNA thermal stability. As such, RNA ligands also serve as fundamental temperature sensors that signal stress through decreased conformational regulation of their protein partners as was observed in these interactome dynamics.

## Introduction

The ability to adapt to changing environmental conditions is among the most fundamental selective advantages enabling evolution of life on this planet. As such, heat shock response (HSR) is a highly conserved complex cellular protective mechanism common to prokaryotes and eukaryotes and provides fitness advantage through the correct folding and/or degradation of proteins after cellular stress and disruption of protein homeostasis. A well-studied step in HSR involves chaperone release of heat shock factor 1 (HSF1) monomers that then activate transcription of many genes involved in HSR, including the stress-inducible heat shock proteins HSP90 and HSP70 (*1, 2*). However, molecular details of heat-induced protein unfolding or features of molecular recognition that precede and mediate HSF1 activation have been more difficult to elucidate. These include alteration of intra- and inter-protein molecular interactions referred to here as interactome changes. Early stages of HSR have been shown to involve only limited changes in protein abundance levels, with most involving decreased protein levels due to increased proteolysis and/or secretion (*3–5*). To gain new insight on interactome changes that occur early in HSR, *in vivo* quantitative protein cross-linking-mass spectrometry (qXL-MS) with isobaric quantitative Protein Interaction Reporter (iqPIR) technology was used to identify significant cellular changes that precede protein abundance and gene expression regulatory pathway changes (*6–8*). These results yield initial insights on time-dependent interactome remodeling and reveal interactome changes that constitute elements of heat stress signal transmission and reception.

## Methods

Information on HeLa cell culture, cross-linking and heat stress of HeLa cells, sample preparation, monomeric avidin enrichment, LC-MS methods for interactome and proteome analysis, proteome and interactome data analysis and luminescent ATP assay are included in **Supplementary materials**.

## Results

For this study, three categories of cross-linker modified lysine-containing peptide species were identified and quantified, including intra- and inter-protein cross-linked and dead-end (DE) peptides where one cross-linker reactive ester group hydrolyzed and only a single lysine site was covalently modified during cross-linking. In total, 5348 non-redundant DE peptides and 5096 non-redundant cross-linked species (3969 intra- and 1127 inter-linked) were identified with <1% FDR from 8 cross-linked samples, derived from 1487 non-redundant proteins. **Figure 1A** shows pairwise correlation plots of log_2_ (42 °C /37 °C) ratios with Pearson correlation (R) coefficient values for all 8 cross-linked samples. Biological replicate samples with the same heat stress duration are well-correlated on the global scale, with R = 0.61, 0.67, 0.88, and 0.77 for 5, 10, 30, and 60 minute heat stress duration, respectively. Summaries of proteome and interactome datasets are included in **Supplementary Table 1** and **2**. A total of 1744 nonredundant cross-linked species were quantified with 95% confidence interval ≤ 0.5, with ≥ 3 ions, and 0 allowed missing values across all heat stress time points. These included 813 intra-linked, 283 inter-linked, and 648 DE peptide species from 433 proteins. A total of 1348 protein levels were quantified with ≥ 3 contributing ions and 0 missing values across all heat stress samples. **Fig. 1B** shows the combined distributions of heat stress-induced proteome and interactome changes at all 4 time points. At the proteome level, no significant changes greater than log_2_ = +/−0.5 were observed at any time point, and ratio distributions for 5-30 minutes appear nearly identical, with slight dispersion at 60 minutes, indicating the lack of significant changes in regulatory protein degradation or production pathways for large numbers of proteins. However, interactome ratio distributions showed larger changes as compared to the proteome data, with increasing number of cross-linked species quantified with log_2_ ratio changes > 0.5 and < −0.5 with increasing duration of applied heat stress. These observations reveal stress-induced alterations of protein conformations and/or protein interactions that occur independently of protein abundance level regulation.

**Figure 1.**
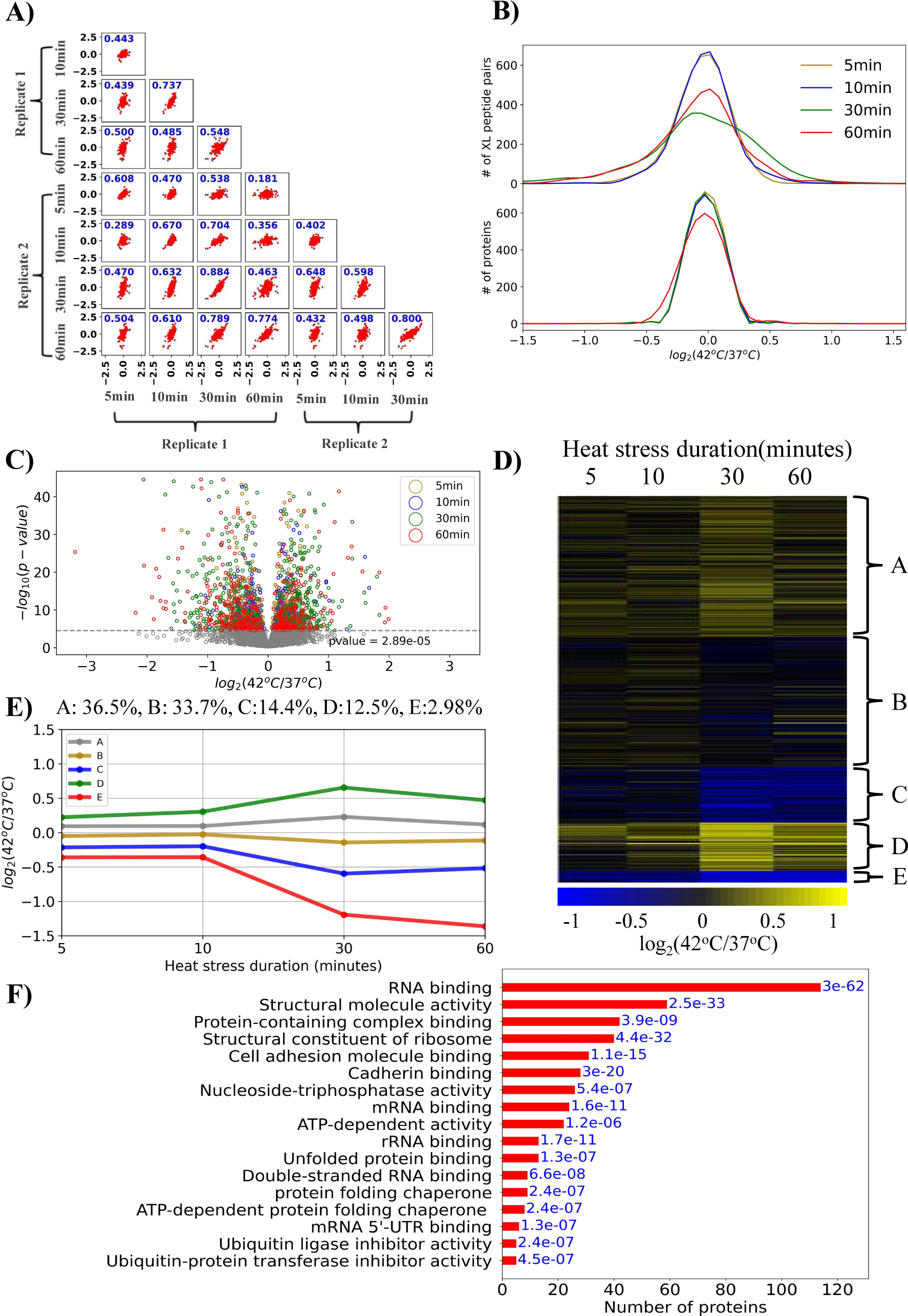
A) Pairwise correlation plots of confidently quantified cross-links (in all replicates and 95% CI < 0.5 for each ratio) with Pearson’s R values for 8 pairs of cross-linked samples. B) Combined distributions of log_2_ values of ratios (42°C/37°C) of confidently cross-linked (XL) peptide pairs, DE peptides and proteins. C) Volcano plot produced with the cross-link log_2_ ratios and calculated P values based on all quantified ions from combined biological replicates. Gray line indicates the Bonferroni-corrected p=0.05 cutoff for significant changes. D) Heatmap and corresponding E) line graph indicating trends and percentages of 5 clusters of cross-linked peptides resulting from K-means clustering of observed cross-link log_2_ ratios. F) Enriched GO molecular function terms and the number of proteins present in cross-links in clusters C and E, and cluster D. Enrichment P-values in blue.

The volcano plot illustrated in **Fig. 1C** was produced with the cross-link log_2_ ratios and calculated P values based on all quantified ions from combined biological replicates and includes Bonferroni correction to identify changes with less than 5% chance of having a true log_2_ ratio of 0 (P < 2.89×10^−5^). The interactome dataset was subjected to k-means cluster analysis (n=5) as shown in **Fig. 1D** to identify distinct groups of cross-linked peptides with similar heat stress-dependent changing patterns (**Fig 1E and Figs S2A-S2D)**. Clusters A and B contain the majority of cross-linked peptide ratios (>70%), with levels largely unaltered during heat stress. Cross-linked peptides in clusters D (13%) and C (14 %), E (3%) display heat stress duration-dependent increasing and decreasing levels, respectively (**Table S3 and S4**). Gene Ontology (GO) term analysis of the 261 proteins with cross-linked peptides in clusters C-E revealed significant enrichment in structural changes among proteins with GO molecular function terms including RNA binding, mRNA binding, rRNA binding and double-stranded RNA binding (**Fig. 1F**). In fact, the single largest group of proteins with cross-linked peptides in clusters C-E (110/223) are associated with the GO term RNA binding. Therefore, quantitative interactome changes reveal significant enrichment of alterations among RNA binding proteins (RBPs) to be intricately linked to early molecular changes in heat stress that occur prior to change in protein abundance levels (**Figs. S2E-S2H**).

## Discussion

### Interactome dynamics during heat stress signal transmission

Previous studies indicate that changes in the RNA-protein interactome can enable cells to respond to both intracellular and extracellular stress (*12, 13*) and UV cross-linking, combined with RNA purification and protein identification revealed changes in RNA-protein interactions with heat stress (*14*). Therefore, one possible explanation of the interactome changes presented here is that heat stress-induced instability of RNA could mediate a loss of intrinsic RNA-binding specificity and affinity (*14–17*) and in doing so, affect the structures of many RBPs as observed here. If so, selective pressures promoting the evolution of RNA-protein interactions, such as beneficial structural stabilization of RBPs, might also include a fitness benefit by sensing and signaling stress through the loss of RBP stabilization, giving rise to the large proportion of RBP interactome changes. This suggests at least some RBPs that lose RNA-mediated structural stability are also client proteins for heat shock chaperone systems that mediate release of HSF1 for downstream HSR activation. RBPs including several helicases were identified as HSP90 client proteins (*18*) and heat stress-induced interactome changes in one such client discussed below appears consistent with the notion of an RNA-RBP stress sensing-signaling system.

Among the RBPs with significant heat-stress induced cross-linked level changes in cells, ribosome proteins account for 33/223 of the cross-linked peptide pairs in clusters C, D and E and the largest increased and decreased cross-link levels observed with 60 minutes heat stress. **Figure 2A** illustrates a heatmap of cross-link levels from ribosome proteins RL35A, RL6, RL5 and RS9. A heatmap of all observed ribosome cross-linked peptide pair levels, 63 in total, is shown in **Figure S3**. Solvent accessible distances for each observed cross-link are illustrated on the ribosome structure (PDB: 7O7Y) and the trajectories or paths are colored by quantitative change observed at 5, 10, 30 and 60 minutes as shown in **Fig. 2B, Fig. S4 and Supplemental Video 1**. Among these, 9 cross-link levels decreased by 2 fold or greater (log_2_ <= −1) from ribosome proteins RL35A, RL6, RL5 and RS9 and one cross-link level increased greater than 2 fold from protein RL35A. The greatest stress-induced change in any cross-link level at 60 minutes was the inter-link RL35A K73-K192 RL6 (Log_2_ = −3.2), while the intra-link between RL35A K73-K45 exhibited the greatest heat stress-induced ribosomal increase (Log_2_ = 1.1) (Fig. 2A). These changes highlight this region of the ribosome large subunit to be among the most heat stress sensitive detected in the present experiment. Integration of DE peptide quantitation revealed that levels of K73 and K192 DE peptides both increased at 60 minutes, indicating increased solvent accessibility/reactivity of these sites at duration of 60 minutes. Therefore, the decreased K73-K192 inter-link level is not resultant from decreased accessibility or reactivity but most likely due to reduction in site proximity to one another. The observed intra- and inter-linked peptides and coordinated quantified DE peptide levels at 60 minutes are shown in **Fig 2C** and S. Video 1. While all ribosome proteins have RNA-contacting surfaces, 28S rRNA interactions appear to nearly surround RL35A and RNA contacts with RL6 are on all sides. Thus, RNA interactions likely play a crucial role in maintaining conformations of both proteins as well as maintaining the RL35A-RL6 ribosomal protein interaction. Therefore, these largest ribosomal interactome changes are consistent with heat stress destabilization of RNA resulting in altered RL35A and RL6 conformational stability as well as the decrease in RL35A-RL6 inter-link levels. Aside from RL35A K73-K45, all other intra-link levels in RL35A (K95-K8, K15-K29, and K95-K29) and in RL6 (K262-K247 and K262-K251) also decrease with heat stress (Fig. 2A), consistent with loss of conformational stability of both proteins in cells exposed to heat stress.

**Figure 2.**
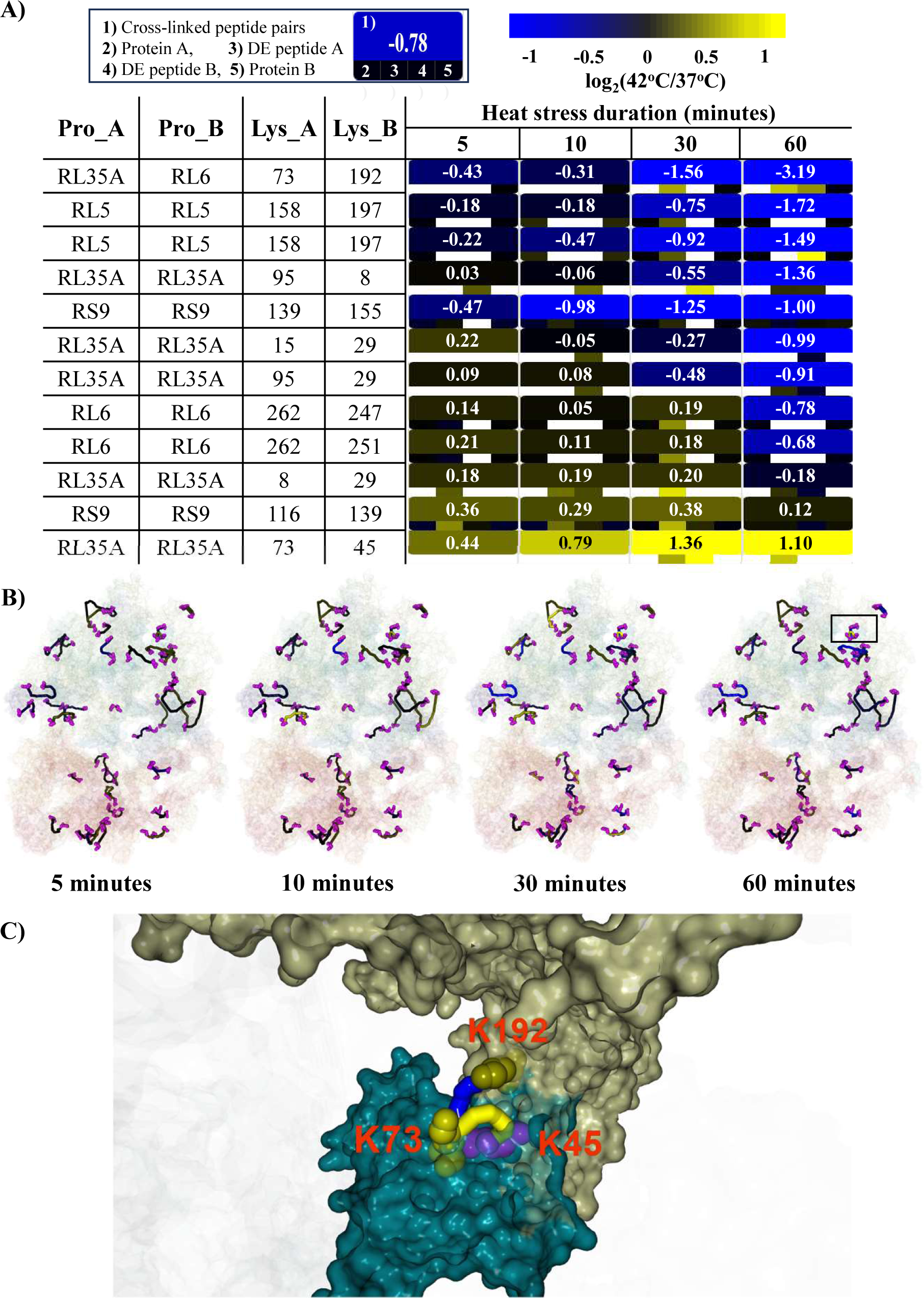
A) Cross-linked and DE peptide levels from largest ribosome changes in RL35A, RL6, RL5, RSX4 and RS9 proteins. Legend (upper left) in this and all subsequent figures indicates map of simultaneous quantitative display of cross-linked, protein and DE peptide levels. B) Jwalk trajectories of all ribosome cross-linked peptides displayed on PDB: 7O7Y colored by quantitative change observed at 5, 10, 30 and 60 minutes. Box indicates the region with the greatest heat stress-induced changes at 60 minutes containing RL35A and RL6. C) RL35A (teal surface) and RL6 (beige surface) illustrating largest ribosome changes observed at 60 minutes in K73-K192 and K73-K45 cross-linked peptide pairs. Space-filled representations of lysine residues K192 (dark yellow) and K73 (yellow) indicate quantified DE peptide levels at 60 minutes. Magenta space-filled residue (K45) indicates no DE peptide quantification at that time point.

As a class, RBPs are generally less well-characterized structurally, in part due to the inherent flexibility of RNA and the conformational changes that occur with binding in both RNA and RBPs (*19*). However, one RBP for which RNA bound and unbound structures are available is Eukaryotic translation initiation factor 4A III (IF4A3) which exhibits remarkable conformational plasticity as illustrated in an RNA-bound structure PDB:2HYI (*20*) and an RNA-free structure inhibited with a middle domain of translation initiation factor 4G (MIF4G domain) of the splicing factor Complexed With Cef1 (CWC22) PDB:2HXY (*21*). **Figures 3A** illustrates the heat map of IF4A3 cross-linked pair and DE peptide ratios with increased heat stress duration at each time point. These results indicate a significant decrease in K321-K152 cross-linked levels were observed at 30 and 60 but not 5 or 10 minutes of heat stress duration. No significant change in IF4A3 protein levels was observed in any duration of heat stress measured in the experiments performed here as shown in Fig. 3A. Thus, the decreased levels of IF4A3 K321-K152 link observed at 30 and 60 minutes can only be due to decreased reactivity of either lysine site (due to solvent accessibility or modification-level changes) or conformational alteration that decreases cross-link formation. The K321-152 cross-link was mapped on the RNA bound and RNA free IF4A3 structures and solvent accessible distances between residues are calculated (24Å, 63Å, (52-58 A for each of 4 chains in 2HXY) respectively) and shown with color indicative of Log_2_(42°C/37°C) change with 60 minutes of heat stress in **Fig. 3B**. The employed iqPIR molecules have a maximum expected linear reach of 38Å based on combined bond lengths (**Fig. S5**). Thus, the solvent accessible distances shown on the closed (2HYI) and open (2HXY) conformations are within and exceed the possible linkable distance, respectively. The ATP-bound closed conformation also localizes R116 and R166 in RecA1 and R316 and K287 in RecA2 domains proximal to one another to facilitate charge interaction with RNA phosphate groups at the intermolecular interface (Fig 3B inset and **Supplemental Video 2**). Decreased stabilization of IF4A3 due to lower ATP levels or RNA thermal instability would be expected to decrease closed conformation contribution to the IF4A3 structural ensemble within cells treated with heat stress, consistent with the observed decrease K152-321 link levels.

**Figure 3.**
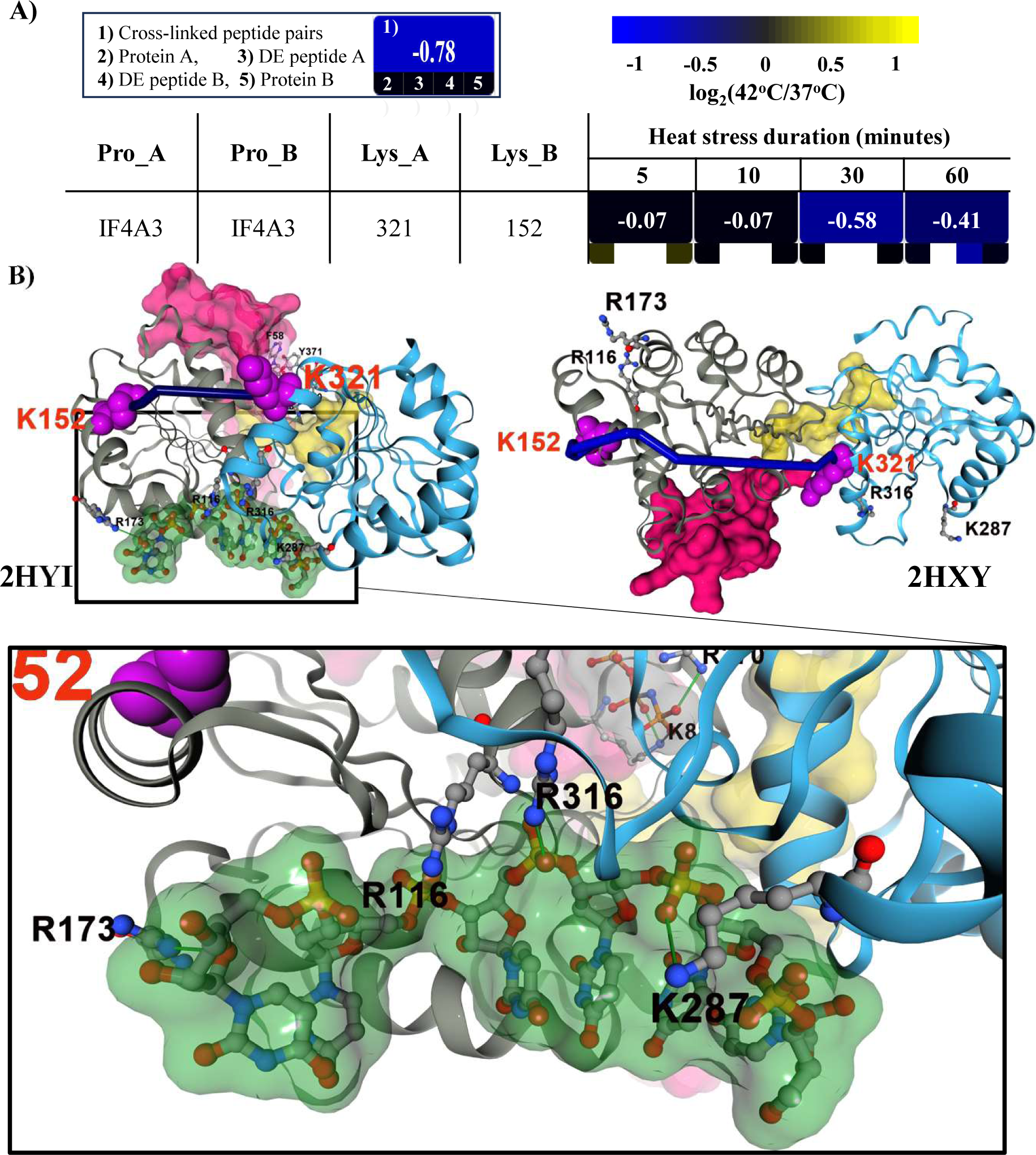
A) Heat map of IF4A3 K321-K152 cross-linked level changes with increasing heat stress duration. B) An RNA-bound structure (2HYI, closed form) and an RNA-free structure (2HXY, open form) of IF4A3. Red surface: N-terminal domain (1–68). Gray cartoon: RecA1 domain (69–239). Yellow surface: flexible linker (240–249). Blue cartoon: RecA2 domain (250–411). Inset illustrates RNA-RBP salt-bridges (green line) that can help stabilize closed conformation.

Like IF4A3, DHX15 is also an ATP-dependent RNA helicase, and has been discovered among many RNA helicases to be a client protein of HSP90 (*18*), requiring this chaperone system for its structural maintenance. With increased heat shock duration, the DHX15 interlink K754-K691 showed a small decrease at all stress duration time points. On the other hand, the K744 DE peptide and K185-K230 cross-link levels showed significant increases and decreases, respectively, as shown in **Fig. 4A**. This DE peptide and cross-link were mapped onto the RNA-bound structure (5LTA) (**Figs. 4B, C and Supplemental video 3**) which showed both appear proximal to RNA interfacial surfaces, indicating this region of DHX15 was increasingly altered with increased stress duration. In addition, K744 reportedly interacts with RNA at the intermolecular interface (*28, 29*). This RNA interaction would decrease reactivity/accessibility of K744 and therefore, increased 744 DE peptide levels with increased heat stress duration are consistent with decreased RNA-DHX15 interactions. On the other hand, the K185-K230 cross-link is within the RecA1 domain and links β4-β5 and β2-α3 loops that are involved in binding RNA and ATP (*28, 30*) by inter-molecular forces. Thus, RNA and ATP interactions stabilize this region in the RecA1 domain and therefore RNA thermal instability would be expected to decrease RNA-mediated conformational stability in this region, altering DHX15 K185-K230 cross-link levels.

**Figure 4.**
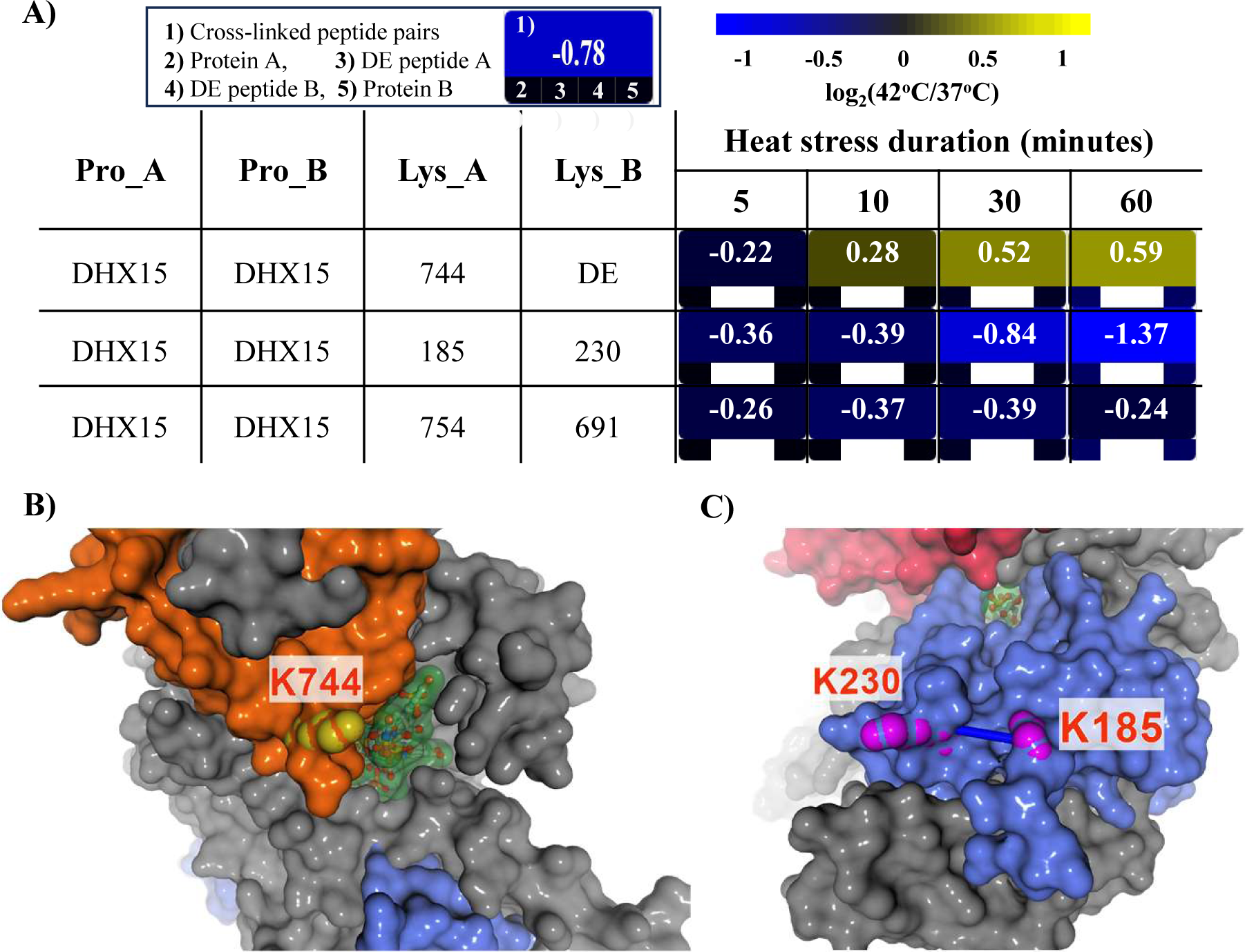
A) Heat map of DHX15 level changes with increasing heat stress duration. B) Site of K744 DE peptide in oligonucleotide-binding (OB)-fold domain (orange surface) that increased with increasing heat stress duration illustrating proximity to RNA (transparent green shaded surface) on PDB: 5LTA. C) Site of K185-K230 link in RecA1 domain (blue surface) on opposite face proximal to RNA binding channel. Gray surface indicates the rest of DHX15 domains (RecA2, ratchet and winged-helix (WH)).

### Interactome dynamics during heat stress signal reception

Unbiased interactome analysis also provides insight on HSP90 molecular chaperone changes that are essential for proteome maintenance, folding, remodeling and maturation of client proteins and modulating HSF1 transcriptional activity (*31, 32*). No significant changes in HSP90 protein levels were observed in any duration of heat stress measured in the present experiments (**Figure 5A)** indicating incomplete HSF1 transcription and translation expected with fully activated HSR. On the other hand, HS90α/ and β cross-linked levels exhibited significant duration-dependent changes (Fig. 5A and **Supplemental Video 4**.), indicating stress duration-dependent signal reception by this master HSF1 regulatory hub. **Figure 5B** shows the ATP ratios (42°C/37°C) determined with luminescence assays based on intracellular ATP levels in heat stressed (42°C) and controlled (37°C) cells with increased heat stress duration. Previous *in vivo* ^31^P NMR studies revealed a sharp decrease in cellular ATP levels of up to 50% in eukaryotic cells within a few minutes of applied heat stress after which ATP levels stabilized at new lower steady state levels (*36*). Luminescence measurements of ATP levels in heat stressed cells performed here also show an initial significant ATP level decrease at 42°C that appears to stabilize at approximately 50% of levels observed in cells maintained at 37°C for the same duration. This heat stress-mediated decrease in ATP levels would be expected to affect partitioning HSP90 conformations among loading, maturation and other conformations as discussed below. HSP90 cross-linked peptide pairs especially informative of conformational change include the homologous links K443-K443 and K435-K435 arising from HSP90α/ and β homodimers, respectively. As shown in **Figures 5C and D**, and S.Video 4, this link spans the HSP90 dimer middle domain (MD) lumenal cavity and would therefore be expected to be sensitive to MD adaptations within the cellular HSP90 structural ensemble of open, client-, ATP-, ADP-, and co-chaperone-bound conformations (*33*). All K443-K443 (α-α)/ K435-K435 (β-β) and K435-K443 (β-α) dimer cross-link log_2_ values all exhibit coordinated trends and appear negative at 5 minutes, but are positive at 10, 30 and 60 minutes of stress duration in cells. Observed cellular HSP90 K443/K435 DE peptide log_2_ values similarly appear negative at 5 minutes of stress, and then are positive at all other time points where quantified. These DE peptide levels indicate that with 5 minutes of cellular heat stress, HSP90 conformations with decreased K443/K435 accessibility/reactivity are enriched at 42 ⁰C compared to those present at 37 ⁰C. With 10 minutes or longer heat stress duration, conformations with increased K443 accessibility are enriched at 42 ⁰C relative to the 37 ⁰C ensemble. The coordination of all K443/K435 DE peptide and dimer link results above indicate dynamic stress duration-dependent changes in the cellular HSP90 structural ensemble, specifically within the lumenal cavity and serve as indicators that stress signal reception indeed has occurred in HSP90 within these cells.

**Figure 5.**
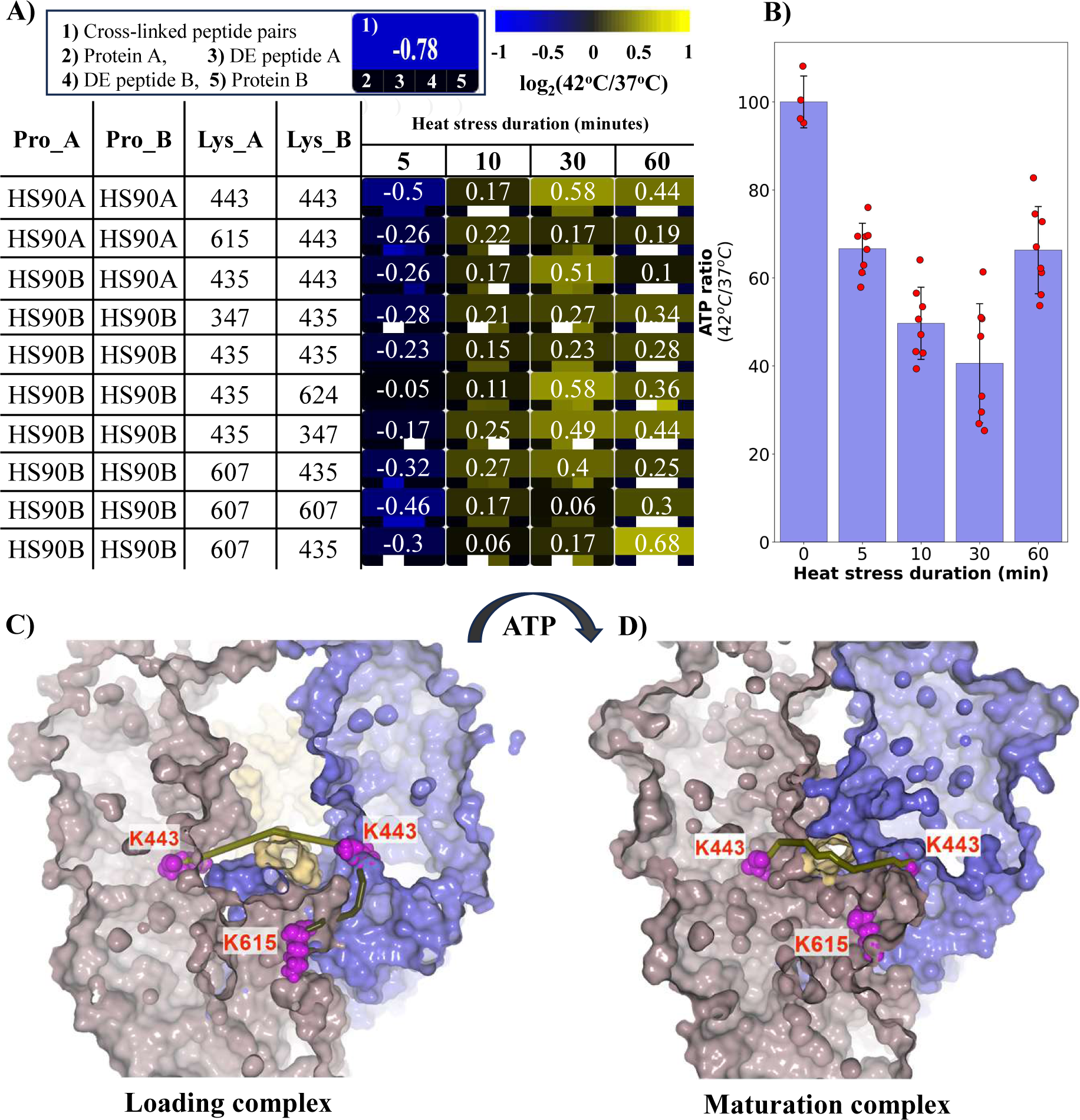
A) Heat map of observed cross-linked HSP90α/β peptide pair, DE peptide and protein levels with increasing heat stress duration. B) ATP ratios (42°C/37°C) calculated by luminescence signals correlated with intracellular ATP levels in heat shocked (42°C) and controlled (37°C) cells with increased heat shock duration. K443-K443 and K443-K615 cross-linked sites illustrated on loading complex C) (left - 7KW7) and maturation complex D) (right - 7KRJ) and reduced K443 accessibility in maturation complex. Solvent accessible trajectory colored to match quantified ratio at 60 minutes heat stress duration.

More complete understanding of these unexpected complex stress-induced HSP90 interactome dynamics will require more comprehensive structural data on HSP90 conformations that exist within the ensemble and additional measurements. Open HSP90 dimers can populate a variety of conformations (*34*) that are currently unresolved, but likely exhibit relatively high K443 accessibilities. Closed HSP90 structures reveal considerable diversity in K443 solvent accessibility levels, and therefore, changing K443 DE peptide and cross-linked levels are indicative of dynamic ensemble compositions with increased stress duration. Among existing human HSP90 dimer structures, the client “loading complex” (PDB:7KW7) exhibits the highest observed K443 solvent accessibility (DSSP calc.=104) and linear/solvent accessible K443-K443 distances between protomers are 34/37 Å, respectively (Figs. 5C and D, and S.Video 4). This loading complex structure also contains HSP70, Hop, the client glucocorticoid receptor and has an empty HSP90 N-terminal domain (NTD) ATP pocket (*34*). Solvent accessibility of K443 in the loading complex is 69% higher than the structure with the next highest accessibility (DSSP calc. = 62), that of the client “maturation complex” (PDB: 7KRJ) (*35*). The maturation complex contains the same client as the loading complex, but differs in the inclusion of p23, does not contain HSP70 or Hop, and contains ATP bound in the HSP90 NTD pocket (*35*). ATP binding triggers NTD lid closure and conformational changes resultant in rate-limiting clamp closure involved in loading-to-maturation complex transition (*34*). As discussed above, heat stress-mediated decrease in cellular ATP levels were observed in these experiments. Therefore, a reduced rate of transition from loading to maturation complexes would also be expected due to rapid reduction in cellular ATP levels after applied heat stress. Taken together, one possible explanation of the observed initial repressed and later increased K443 DE peptide and cross-linked levels involves the combined effects of increased demand for client structural maintenance and subsequent ATP level reduction. Both factors would be expected to affect partitioning of open dimer, loading, maturation and other conformations (**Figure S6A**). Since demand for client structural maintenance is expected to remain high throughout the stress duration, the proportion of open dimer at 42 °C is expected to be lower than that at 37 °C at all time points. At early time points with high ATP levels, relatively fast transition from loading to maturation complexes (and subsequent conformations) is expected (**Fig. S6B**), resulting in lower ensemble contribution of conformations with high K443 accessibility (open and client loading complex) than those of low K433 accessibility (maturation complex and possibly others) at 42 ⁰C compared to 37 ⁰C, resulting in observed negative log_2_ DE peptide and cross-linked ratio values. However, subsequent reduction of cellular ATP levels may increase relative proportion of loading complex conformations since transition at this checkpoint is ATP-dependent and therefore, could give rise to increasing levels of K443, DE peptide and cross-linked levels at longer heat stress duration **(Fig. S6C)**. This is only one of many possible explanations of the observed interactome dynamics and many additional future experiments, structures and efforts for interpretation are required to better comprehend this new type of time-dependent data. Nonetheless, while comprehension of these observed changes may as yet be incomplete, these results demonstrate that qXL-MS can enable visualization of molecular-level interactome changes in cells exposed to even as few as 5 minutes of heat stress, in the absence of any measured proteome level changes. Moreover, these dynamic interactome data reveal elements of both, heat stress signal transmission by many proteins, at least some of which are known clients of HSP90 and heat stress signal reception by this master regulator of heat shock response. Quantitative interactome studies provide exciting new opportunities to further visualize mechanistic details of this and many other processes in cells.

## Conclusion

In living systems, functional regulation is the cumulative product of multifactorial inputs, making molecular details difficult to decipher. A primary conclusion derived from the present work is that the combination of quantitative proteome, interactome and lysine solvent accessibility data sets can enable visualization of time-dependent changes in the ensemble of protein conformations and protein-protein interactions within cells exposed to stimulus. While only a small snapshot of possible time-dependent interactome changes have been recorded in the work reported here, these efforts nonetheless revealed that reproducible and statistically-significant time-dependent interactome changes are detectable in cells exposed to 5⁰C increased temperature, even in periods as short as 5 minutes. Further measurements such as these can likely enable new types of large-scale computational models that can help further increase systems-level understanding of complex cellular pathways involved in heat stress response. Moreover, the examples discussed all involve interactome changes that occur independently of proteome-level adaptation and suggest that in principle, this approach can generally be applied to visualize time-dependent changes in molecular mechanisms involved in many other cellular pathways, such as response to nutrient deprivation, drug or toxin exposure, or virtually any change in cell growth conditions.

While interactome changes in the heat shock protein HSP90 were an expected finding with applied cellular heat stress, the surprising observation of stress duration-dependent HSP90 dynamics indicate the occurrence of additional molecular changes with increasing heat stress duration. Additionally, the integration of DE peptide quantitation provides insight on time-dependent lysine site solvent accessibility/reactivity changes that occur inside cells with increasing stress duration. These changes improve understanding of increased HSP90 inter-link levels with increased stress duration since together they indicate a duration-dependent increase in closed conformations that have increased K443 solvent accessibility. This observation suggests an enrichment in client loading complex conformations since among known structures, that conformation exhibits the highest K443 solvent accessibility. While these aspects of HSP90 dynamics cannot be conclusively determined with this initial time-dependent interactome dataset, previously reported heat stress-dependent decrease in cellular ATP levels and the expected enrichment of loading complexes due to reduced ATP levels seems likely. In any case, the quantitative interactome approach also unexpectedly revealed that the single largest significant enrichment in molecular interaction adaptations during heat stress was not among heat shock proteins, but rather among RNA binding proteins. These interactome changes support a concept of RNA thermal instability as a sensory component of cellular heat stress detection, with disruption of RNA-mediated RBP conformational stability and protein interactions serving in a stress signal messaging capacity. Thus, an additional dimension of fitness benefit provided by RNA-protein interactions may be derived through sensing and signaling heat stress by these complexes. These observations together with the recent discovery that several RNA helicases are themselves, HSP90 client proteins, including DHX15 provide a plausible link between thermally-induced RNA-mediated RBP conformational signaling, increased HSP90 demand and HSF1 activation. Finally, as is true for any new source of quantitative data, the ability to comprehend meaning in the observed changes constitutes the next level of challenge arising with the realization of the ability to acquire the data. Future more widespread interactome dynamic studies with additional applied stress conditions, inhibitors, or other perturbations will help further increase understanding of molecular changes inside cells that can be visualized through quantitative interactomics.

## Data Availability

Mass spectrometry data have been deposited to the ProteomeXchange Consortium via the PRIDE repository under identifiers PXD050165.

Cross-linking data have been uploaded to the XLinkdb online database (http://xlinkdb.gs.washington.edu/xlinkdb/PHP_SAVED_VIEWS/sgp20/Time_Dependent_Heat_Shock_response_Combined.php).

## ACKNOWLEDGEMENTS

The authors acknowledge and thank all members of the Bruce lab for helpful comments and suggestions in the course of this work and manuscript preparation.

## Author contributions

J.B. conceived of the project. S.P. cultured cells, performed time-dependent heat stress, cross-linking and quantitative interactome experiments. N.K. performed quantitative proteome experiments. A.K. developed and applied informatics used for quantitative interactome analyses and comparisons. S.P. and J.B. designed research, prepared the manuscript and videos with input from all authors and lab members.

## Methods Details

### HeLa Cell Cultivation

HeLa cells were grown in 150 mm plates with 37 ⁰C with 5% CO_2_ and a humidified atmosphere with Dulbecco’s modified Eagle media (DMEM), 10% dialyzed Fetal bovine serum (Valley Biomedical) and 100U/mL penicillin-streptomycin (Fischer Scientific). Cells were harvested at approximately 90% confluence. The harvesting involved the aspiration of growth media and a 10 mL wash of PBS, followed by cell detachment with the addition of 10 mL of 20 mM EDTA in a PBS buffer and incubation at 37 ⁰C for 3 minutes. Cells were carefully washed off the plate and collected into 50 mL tubes. Cells were then pelleted at 300 g for 3 minutes and supernatant aspirated. To replace the ions chelated by EDTA, the cells were washed with 10 mL of 1 mM CaCl_2_ and 1 mM MgCl_2_ in PBS, pelleted (as described), and aspirated. This cell washing was performed twice. After the last cell aspiration, a 50:50 (by volume) mixture was made with the cell pellet and the cross-linking reaction buffer (170 mM Na_2_HPO_4_ at pH 8.0). The cells in the 50 mL tube were transferred into eight 2 mL tubes; four for control (37 °C) and four for heat stressed (42 °C) exposed for different time periods (5, 10, 30 and 60 min) as shown in **Figure S1a and b**.

### Cross-linking and heat stress of HeLa cells

The mild heat shock procedure was similar to that described in Mahat et al.(*1*), and is illustrated in **Fig. S1B**. Tubes (2mL) with cells suspended in cross-linking buffer (50/50 (V/V)) were shaken at 650 rpm for 30 minutes at 37 ⁰C. At 30 minutes, the cells in all tubes were pelleted and the solvent aspirated, then equal volumes of preheated cross-linking buffer at 37 ⁰C or 42 ⁰C were added to the pelleted cells in each tube. After that, the cells with cross-linking buffer preheated at 37 ⁰C were maintained at 37 ⁰C and shaken at 650 rpm, and the cells with cross-linking buffer preheated at 42 ⁰C were maintained at 42 ⁰C and shaken at 650 rpm to induce heat stress. Two aliquots of cells one at 37 ⁰C and the other at 42 °C were cross-linked at time points 5, 10, 30 and 60 min.) with iqPIR reagents, each at final concentration of 10 mM. RH iqPIR cross-linker was added to control samples (37 ⁰C) and SH iqPIR cross-linker was added to heat stressed samples (42 ⁰C). Prior to adding cross-linker to control and heat stressed samples, a 50µL aliquot of cell suspension was taken from each tube and cells were immediately lysed for data-dependent acquisition (DIA)-based label-free proteome quantification. Detailed information about the sample preparation for the DIA-based label-free protein quantification is described below. After 30 minutes of cross-linking, all reactions were quenched with an equal volume of 150 mM ammonium bicarbonate. After that, the cells were pelleted and washed three times with 10 mL of 150 mM ammonium bicarbonate to remove unreacted excess crosslinker in solution.

### Sample preparation

The proteome and interactome (cross-linked) cells were lysed with the addition of 10 M urea with 150 mM ammonium bicarbonate in a 60:40 ratio with cells to a final urea concentration of 6 M. The solutions were sonicated with a GE-130 ultrasonic processor for 3 x 5 second bursts running at 50 watts to ensure near complete lysis. Reduction via a 30 min 5 mM TCEP incubation at RT followed by alkylation with 10 mM iodoacetamide for 30 min was performed. Total protein concentration was quantified via a standard Bradford Assay for each sample. Interactome samples from cells exposed to 37 °C or 42 °C were mixed pairwise in 1:1 ratio based on equal amount of total protein (**Fig. S3C**). The total protein quantity of the mixed samples was 2 mg. Interactome and proteome samples were digested overnight (16 hours) with trypsin at a ratio of 1:200 (enzyme : protein) by weight at 37 ⁰C, acidified to a final volume of 1% trifluoracetic acid (TFA) to quench the digestion and prepare the digest for desalting. A 3cc Waters Sep-Pak C18 column was washed with 3 mL of methanol, followed by 3 × 3 mL washing with 0.1% TFA in acetonitrile (ACN), and then 3 × 3 mL of 0.1% TFA in water to equilibrate the column. The sample was added to the column and bound peptides were washed 3 x 3 mL with 0.1% TFA, then eluted in 1mL of 80% ACN with 0.1% formic acid. To construct a chromatogram library for DIA-based proteome quantification, a 200 µL aliquot taken from each 1mL of eluate with Sep-Pak for the label-free digested cells was pooled as described elsewhere (*2, 3*). Samples were dried with vacuum centrifugation. After that, the dried label-free samples were stored in −80 °C freezer for DIA-based proteome quantitation with Thermo Q-Exactive Plus. Interactome samples were reconstituted with 0.5 mL in the strong cation exchange (SCX) buffer A (7 mM KH_2_PO_4_, pH 2.8, 30% ACN) to further reduce unreacted and hydrolyzed cross-linker levels. The entirety of each digested interactome sample was loaded onto the SCX column coupled with an Agilent 1200 series HPLC with an increasing salt gradient. The SCX column used was a Phenomenex Luna 5 µm with 100Å particles in a 250 x 10.0 mm column housing. Specifically, the gradient flowed at 1.5 mL/min going from 0% B (Buffer B = Buffer A + 350 mM KCl) to 5% B over 7.5 minutes to 60% B over 40 minutes to 100%B over 20 minutes, holding there for 10 minutes, then back to 0% B after the washing step for an additional 20 minutes. Fractions were collected every 5 minutes starting at 17.5 minutes. Early and late SCX fractions were pooled to result in 6 SCX-separate samples as 1-5, 6-7, 8, 9, 10, and 11-14, and each was dried by vacuum centrifugation. **Fig. S1D** shows a SCX chromatogram, which was obtained with a cross-linked sample acquired after at 5 minutes of exposure to 42 or 37 ⁰C. For biotin-avidin affinity enrichment of cross-linked species, the pooled fractions were reconstituted with 0.5 mL of 150 mM ammonium bicarbonate (pH 8.0) except for the pooled fraction 11-14. Due to the higher amount of salt than other factions, the pooled fraction 11-14 were reconstituted into 2 mL of 150 mM ammonium bicarbonate. The pH of each pooled fractions was adjusted to 8.0 with 3.0 M NaOH solution to ensure adequate binding and affinity enrichment with monomeric avidin (ThermoFisher Scientific 20228). The final volume after adjusting pH was 0.8 mL for the pooled fractions 1-5 and 6-7, 0.6 mL for the fractions 8, 9, 10, and 2.6 mL for the pooled fraction 11-14.

### Monomeric avidin column methodology

An affinity enrichment with monomeric avidin was performed with an avidin column that was prepared by filling monomeric avidin (ThermoFisher Scientific 20228) slurry into a 100 × 4.6 mm empty column with a 500 mL syringe (**Fig. S1E**). The avidin column was installed on an AKTA pure chromatography 25 (GE Healthcare, Marlborough, MA USA). The column volume (CV) was 1.8 mL. During affinity chromatography, absorbance was monitored at 214 and 498 nm wavelengths. The injection volume was 0.8 mL for fractions 1-5 and 6-7, and 0.6 mL for the fractions 8, 9, and 10. The pooled fraction 11-14 was split into two equal volumes and two injections, each of 1.3 mL were used. The sample was loaded onto the column at a flow rate of 3 ml/min in 150 mM ammonium bicarbonate. After loading the sample, the column was washed with 3 CV of 150 mM ammonium bicarbonate, and then switched to a 70% ACN and 30% water solution containing 0.1% TFA for elution of cross-linked peptides with a total of 12 CV. Elution peaks from the avidin column were collected and dried under vacuum.

### LC-MS methodology for interactome analysis

After monomeric avidin enrichment, the dried samples were reconstituted in a 2% ACN and 98% water solution containing 0.1% FA. All fractions were analyzed two times each with 4.0 µL of sample injected onto a Waters nanoAquity UPLC (nanoACQUITY, Waters, Manchester, UK) coupled to a Thermo Q-Exactive Plus instrument (**Fig. S1F**). For cross-linked peptide analysis, the nanoAquity UPLC was equipped with a 60 cm C8 column, and a 5 cm trap column, both packed in house following an established procedure (*4*) with 5 µm Reprosil C8 with 120Å pores into 75 µm ID fused silica columns. The sample was loaded onto the trapping column at a flow rate of 2 µL per minute for 10 minutes in 100% buffer A. After trapping, the mobile phase gradient was applied as follows: 2% B to 18% B over 3 minutes, then forming a shallow gradient from 18% to 40% B for 118 minutes (totaling 121 minutes since loading), followed by 80% B for 20 minutes then 2% B for 20 minutes for column equilibration. A flow rate to sperate peptides was 0.3 µL/min. The mass spectrometer employed a top 5 data dependent method, selecting for 4^+^ to 7^+^ ions to maximize selection of cross-linked peptides. MS1 and MS2 scans were both run with 1 µscan at 70k resolution with an AGC of 1e6 and 5e4, respectfully. The spray voltage was run between 2.3 - 2.7 kV. All samples were run this way except for a pooled sample of fractions 1-5, where the charge exclusion was lowered to 3^+^ to 7^+^ to allow for more dead-end peptide identifications. HCD normalized collision energy (NCE) was set to 30 with a 3 m/z isolation window and dynamic exclusion of 30 seconds.

### LC-MS methodology for proteome analysis

For DIA-based label-free peptide analysis, peptides that were taken from each control and heat stress samples prior to adding cross-linker as described above were separated with a 30 cm x 75 µm ID C18 (5 µm Reprosil C18 with 120Å pores) column, and a 5 cm trap column on a nanoAquity UPLC. Samples were loaded onto the trapping column at a flow rate of 2 µL per minute for 10 minutes in 100% buffer A. After trapping, the peptides were separated at 0.3µL/min along with linear gradient condition; 5% B to 35% B over 90 minutes, 35% to 60% B for 10 minutes, and 60% B to 95% B for 5minutes, followed by 2% B for 20 minutes for column equilibration. Mass spectrometer setup for DIA was based on that described elsewhere (*2, 3*). The sample for generating the chromatogram library was resuspended with 20µL of 0.1% formic acid in 20% ACN and 80% water for loading it to nanoAquity UPLC coupled to Thermo Q-Exactive Plus and analyzed by injection of 1.0 µL of the sample. The chromatogram library was constructed by six DIA spectra acquired by an overlapping window pattern from narrow mass ranges with 4 m/z isolation window at 17,500 resolution, 1×10^6^ automatic gain control (AGC) target, 55 milliseconds (ms) of maximum inject time, 27 collision energy (CE) and 27 loop count. Two precursor spectra, wide mass spectrum (400 - 1600 m/z) and narrow spectrum (395 - 505, 495 - 605, 595 - 705, 695 - 805, 795 - 905, and 895 - 1005 m/z), were acquired with 35,000 of resolution, 3×10^6^ of AGC target, and 100 ms of maximum inject time. For quantitative analysis of each sample, the mass spectrometer was set up with 385 - 1015 m/z of mass range at 35,000 resolutions, 3×10^6^ of AGC target and 100 ms of maximum injection time for precursor spectra, 24 m/z of precursor isolation windows at 35,000 resolutions, 1×10^6^ of AGC target, 55 ms of maximum inject time and 27 of NCE for DIA.

### Proteome analysis

The analysis of DIA data was started with converting the acquired DIA mass spectra to .mzML files using MSConvert (version 1.10.19) (*2, 3*). After that, a chromatogram library for analyzing the samples was generated by Walnut in EncyclopeDIA (*2, 3*) with the default settings: fixed cysteine carbamidomethylation, 10 ppm of precursor and fragment tolerances, using Y ions, and 1 maximum missed cleavage. Library searching in EncyclopeDIA also was performed with the default settings; 10 ppm of precursor and fragment tolerances, using B and Y ions, and 10 ppm of library mass tolerance. The results were filtered to a 1% protein-level false discovery rate (FDR), followed by a 1% peptide-level using Percolator 3.1 (*2, 3*) during library searching with EncyclopeDIA.

### Interactome analysis

LC-MS .Raw files were converted to mzXML format prior to searching for PIR relationships with the Mango algorithm(*5*). The MS2 Mango generated files were then searched with Comet search engine(*6*) against the Uniprot obtained proteome database for *Homo sapiens* (*7*). The Comet search generate pep.XML files and analyzed with XLinkProphet(*8*). The resulting cross-link identifications were then filtered to an FDR < 1%. Data analysis steps for ion apportionment and quantitation to derive cross-linked peptide ratios and p values were performed as previously described(*9*). The reporter ions were not quantified to eliminate any contributions to the signal that may result from co-isolation of other cross-linked precursor ions (**Fig. S1G**). Because of this, all quantification was performed with peptide and fragment ions containing the released portion of the iqPIR reagent. Once all peptide fragment contributions were assigned, a global median normalization factor was applied to adjust the log_2_ ratios. Fragment quantifications were then combined per identified peptide with an additional outlier removal procedure to obtain more accurate and reproducible quantifications. After cross-linked peptides were quantified in all datasets, ratios were averaged statistical filters were applied to identify significant changes. The biological replicates of each sample were merged with applied statistical filters to create the combined distribution. Data analysis was performed with the combined data set with 95% confidence intervals less than or equal to 0.5, each quantification required at least 3 independently quantified ions and maximum one missing value across all time points. The filtered data was then grouped into 5 clusters with K-means algorithm regarding log2 ratios. The entire interactome dataset with quantification, structures with mapped cross-links and k-means clustering assignments are available to view online in XLinkDB and supplement data. An ontology enrichment analysis, correlation plots between biological replicates, log2ratio distribution plots, volcano plots were performed with the data set downloaded from XLinkDB (http://xlinkdb.gs.washington.edu/xlinkdb/PHP_SAVED_VIEWS/sgp20/Time_Dependent_Heat_Shock_response_Combined.php). The ontology enrichment analysis was performed with the proteins corresponding to cross-links including inter, intra, homodimer and deadend that exhibited large statistical changes. The comparison of log2ratio distribution was performed with the overlapping proteins represented in both cross-linked and label-free samples. Solvent accessible distances for each observed cross-link were calculated with Jwalk (*10*). DSSP solvent accessibilities were calculated for all lysine sites within XLinkDB (*11*).

### ATP luminescent assay

Cellular ATP levels with different heat shock duration were measured using CellTiter-Glo luminescent cell viability assay kit (Promega, G7570) following the manufacturer’s instructions with a little modification(*12–15*). Briefly, the pelleted cell harvested as described in the HeLa cell cultivation section above was resuspended with a cross-linking reaction buffer and 50 µL aliquot of the resuspended cells was transferred into thirty-six 2 mL tubes for control (37 °C) and heat stressed (42 °C) for different time periods (0, 5, 10, 30 and 60 min). All 2 mL tubes with cells suspended in cross-linking buffer were shaken at 650 rpm for 30 minutes at 37 ⁰C. At 30 minutes, the cells in sixteen 2 mL tubes were maintained at 37 ⁰C, and the cells in another sixteen 2 mL tubes were transferred to a 42 ⁰C heat block to induce heat stress. The cells in the rest of 2 mL tubes (four tubes) were pelleted and the cross-linking buffer aspirated, then 100 µL of DMEM was added to the pelleted cells in each tube to measure ATP level at 0 min time point. The cells resuspended with DMEM were transferred to a 96-well plate and then added 100 µL of CellTiter-Glo reagent to the cells. After that, the plate was placed on a plate shaker for two minutes to induce lysis and allowed to incubate at room temperature for 10 minutes prior to measuring luminescence using the Cytation 5 cell imaging multi-mode reader (BioTek). At each time point (5, 10, 30 and 60 min), the cells in the eight tubes four from control (37 °C) and the other from heat stressed (42 °C) were palleted, resuspended with DMEM and added 100 µL of CellTiter-Glo reagent to the cells to measure luminescence like the cells prepared for 0 min time point as described above.

## Discussion

### Ribosome proteins

From all structures currently available in the Protein Databank, the existing ribosome structure PDB:7O7Y was found to map to the largest number of cross-linked peptide pairs (316) from the entire time-dependent heat stress dataset presented here.

### Heat shock proteins

Heat shock response can also be induced by cellular treatment with HSP90 inhibitors such as 17AAG in the absence of any applied heat stress (*16*). This is thought to occur because 17AAG can disrupt a conformation-specific HSP90-HSF1 interaction, prolonging HSF1 occupancy of the HSP70 promoter and increasing HSF1 transcriptional activity of heat shock proteins (*17*). Thus,heat shock response due to 17AAG treatment would be expected to be independent of the proposed RNA-RBP heat stress sensing and signaling mechanisms and therefore, would not be expected to include changes in RBPs such as RL35A-RL6 or IF4A3 cross-linked peptide levels discussed above. Indeed, previous interactome data collected with N-terminus inhibitors, including 17AAG, Ganetespib and XL-888, of MCF-7 cells indicated heat shock response activation of HSP90α with increased protein and cross-link levels. However, the RL35A-RL6 K73-K192 and IF4A3 K152-K321 cross-link levels were unaltered with cellular 17AAG, Ganetespib or XL-888 treatments (*18*) (**Figure S7**). These combined observations indicate that the significant heat stress-induced changes observed in RL35A, RL6 and IF4A3 are independent of mechanisms downstream of HSF1 transcriptional activation, consistent with notion of RNA-RBP sensing and signaling early on during heat stress prior to increased HSF1 release.

### DHX15 protein

In their efforts to identify helicases as HSP90 clients, DHX15 protein levels were observed to be decreased 1.6 and 1.4-fold with HSP90 inhibition with Onalespib and Alvespimycin inhibitors, respectively, indicating that like many other HSP90 clients identified in their work, DHX15 requires active HSP90 for folding and structural maintenance (*19*).

---

**Figure S1.**
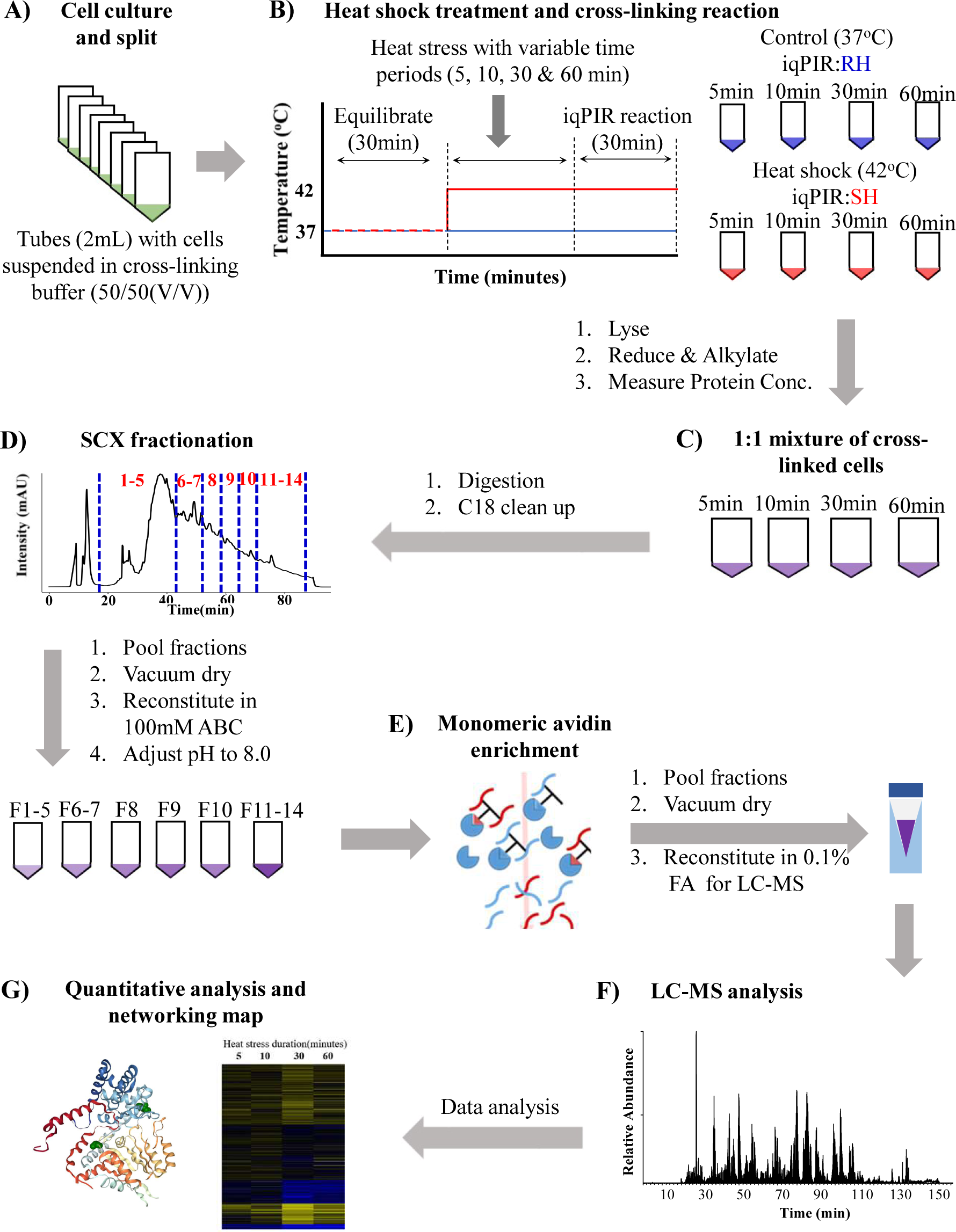
A) Cell suspension split to 2mL tubes after cell culturing in a 50mL. B) Cross-linking of the cell suspension with iqPIR reagents (37 ⁰C = RH, 42 ⁰C = SH) after heat shock for different time periods (5, 10, 30 and 60min). C) 1:1 mixture of cross-linked cells based on their protein concentrations after lysis, reduction and alkylation. D) SCX fractionation after sample digestion and C18 clean up. E) Avidin enrichment. F) LC-MS analysis. G) Data analysis with Comet, Mango, and XLinkProphet to identify crosslinks.

**Figure S2.**
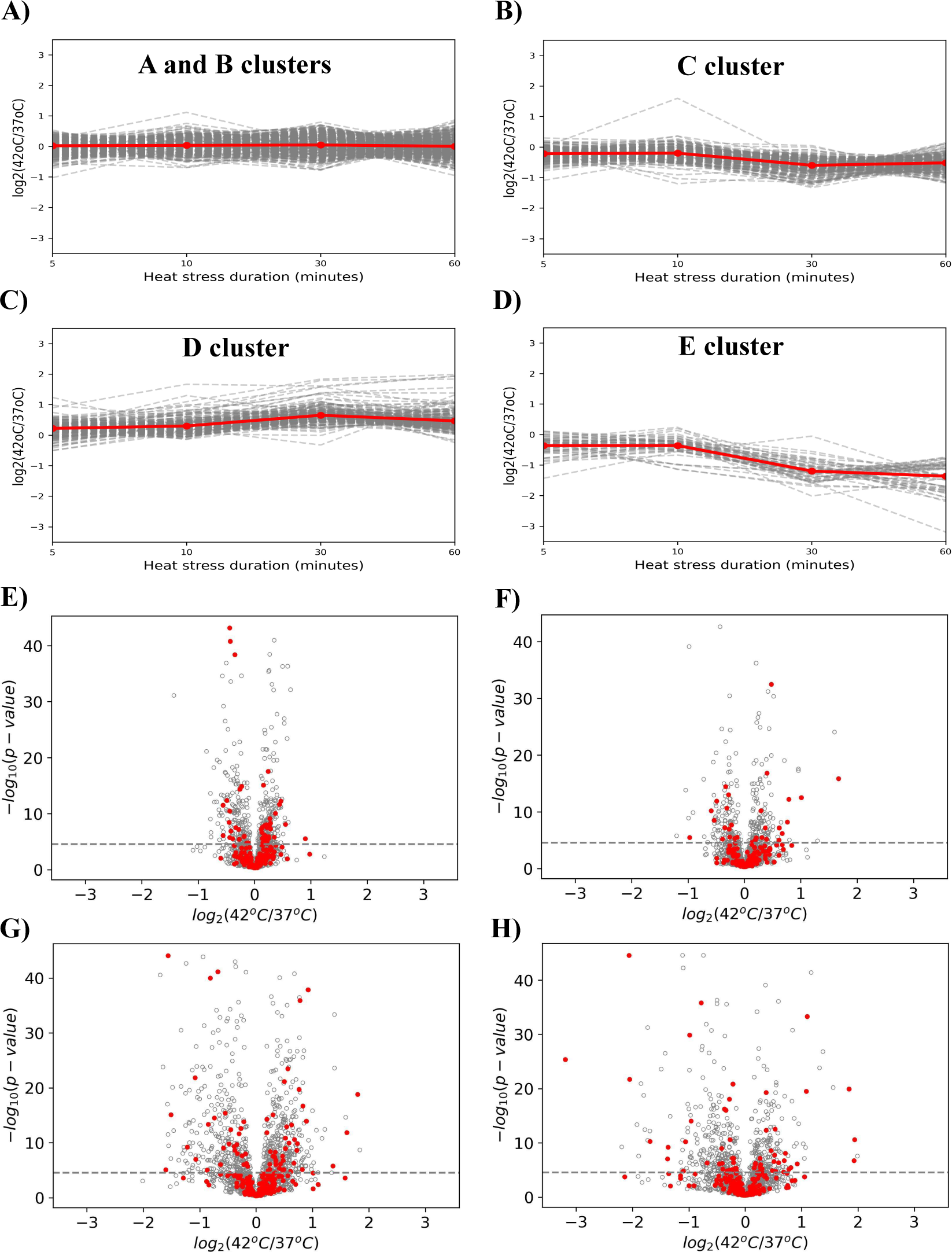
Line plots indicating distinct groups of cross-linked peptide pairs with heat-stress-dependent changing patterns resultant from K-means clustering analysis. A) A and B clusters, B) C cluster, C) D cluster and D) E cluster. The red lines correspond to the mean of trajectories in the same cluster. Volcano plots showing RNA binding proteins highlighted with red color. E) 5min, F) 10min, G) 30min and H) 60min.

**Figure S3.**
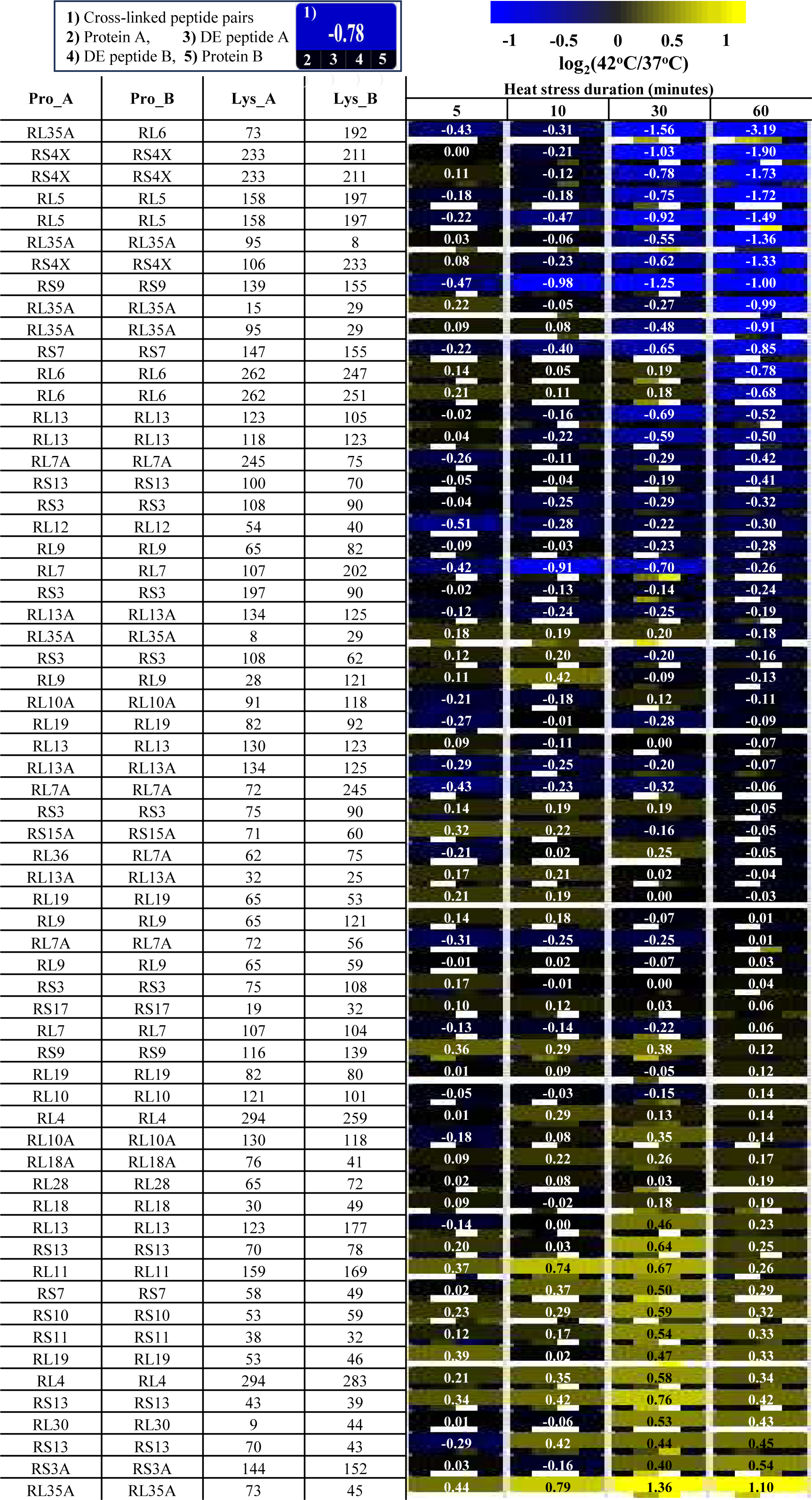
Heat map of all observed ribosome cross-linked peptide pairs.

**Figure S4.**
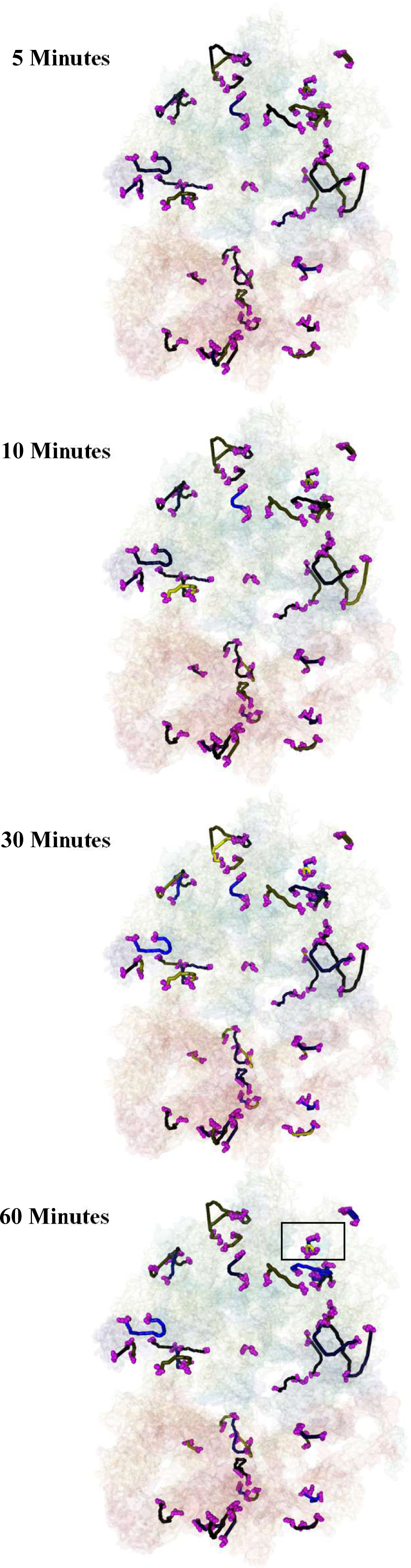
Jwalk trajectories of all links on 7O7Y colored by quantitative change observed at 5, 10, 30 and 60 minutes. Box indicates region with greatest heat stress-induced changes containing RL35A and RL6.

**Figure S5.**
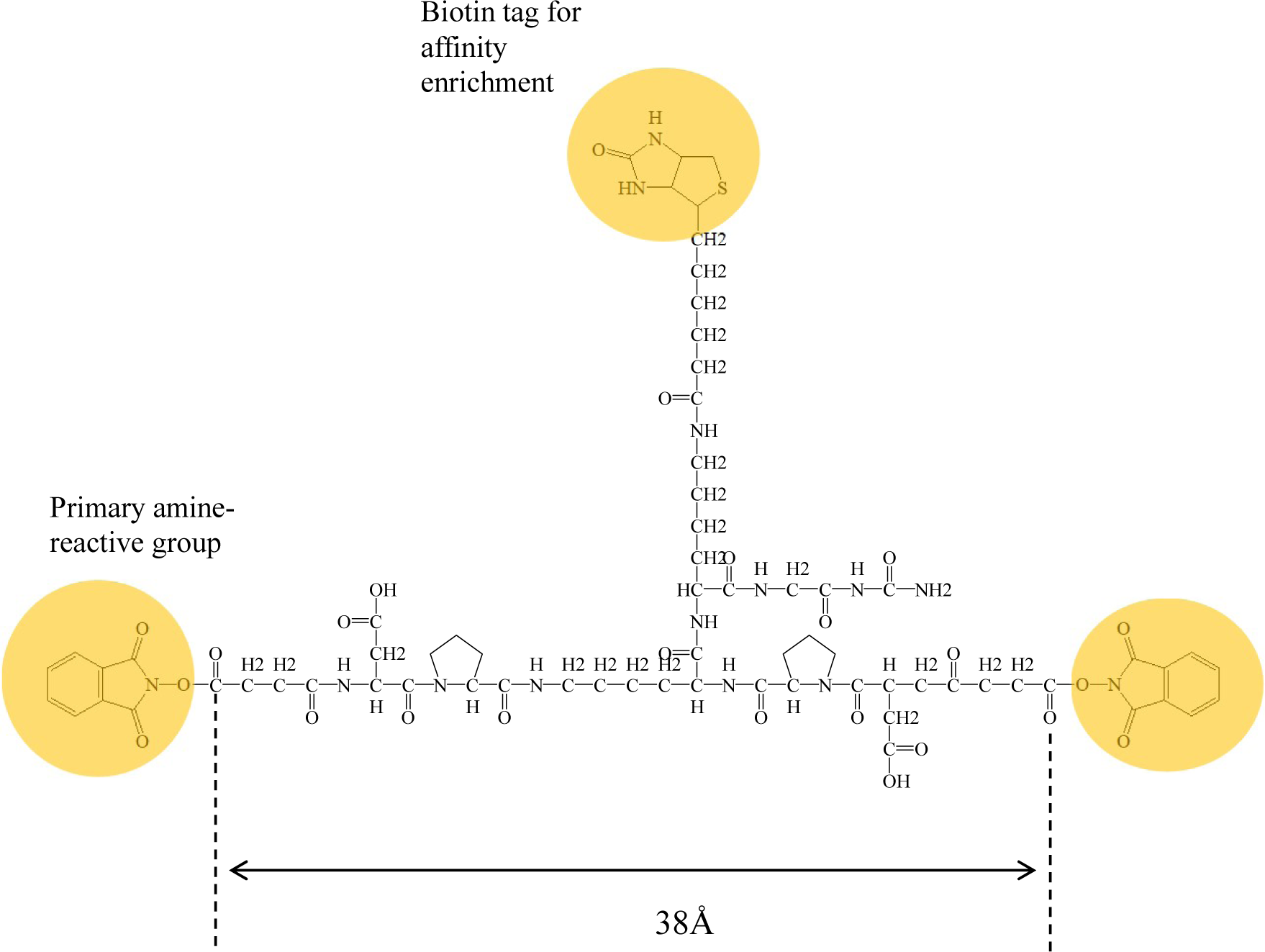
An iqPIR cross-liner structure and a maximum expected linear distance between two reactive group.

**Figure S6.**
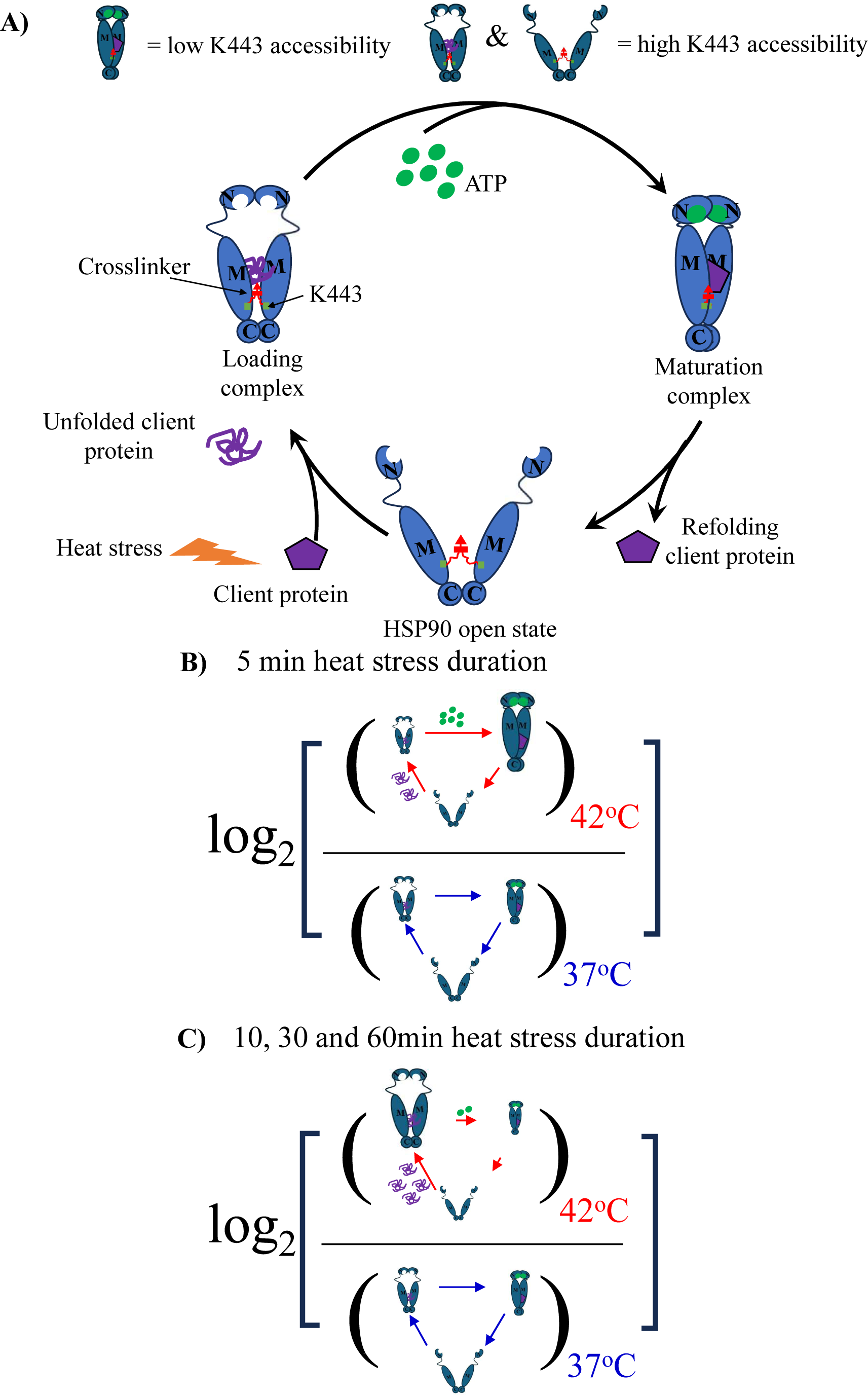
A) HSP90 chaperon cycle. After loading unfolding client protein to an open state, HSP90 undergoes a conformational change to a loading complex state. ATP binding to the N-terminal domain induces formation of a compact APT-bound closed state (maturation complex state). HSP90 returns to the open state with APT hydrolysis. In this cycle, the open and loading complex states have a higher K443 accessibility than that of the maturation complex state. The size of each conformational state and the number of green spots (ATP) in log_2_ equations (B) and C)) represent the expected proportions of each state and ATP levels, respectively, at different heat stress duration.

**Figure S7.**
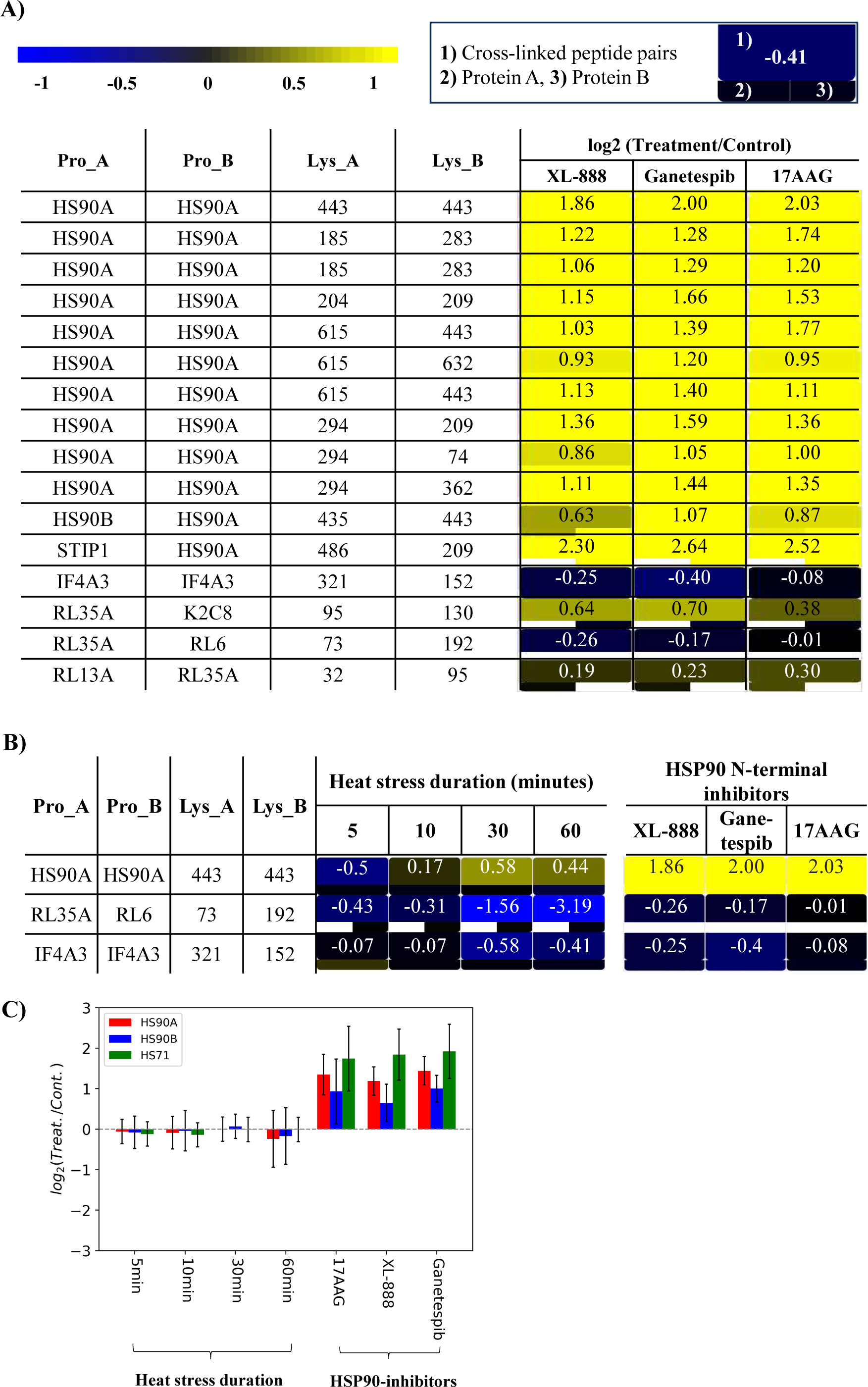
A) Heat map for observed HSP90, RL35A, RL6 and IF4A3 proteins with XL-888, Ganetespib and 17AAG treatment of MCF-7 cells. B) Comparison of cross-linked peptide pair and protein level changes stimulated by heat stress and HSP90 N-terminal inhibitors. Bar graph showing protein level changes with heat stress and HSP90 inhibitors.

## References

1. I. Shamovsky, M. Ivannikov, E. S. Kandel, D. Gershon, E. Nudler, RNA-mediated response to heat shock in mammalian cells. Nature 440, 556–560 (2006).

2. Y. Chin et al., Targeting HSF1 for cancer treatment: mechanisms and inhibitor development. Theranostics 13, 2281–2300 (2023).

3. V. Cappelletti et al., Dynamic 3D proteomes reveal protein functional alterations at high resolution in situ. Cell 184, 545–559.e522 (2021).

4. A. Finka, V. Sood, M. Quadroni, L. Rios Pde, P. Goloubinoff, Quantitative proteomics of heat-treated human cells show an across-the-board mild depletion of housekeeping proteins to massively accumulate few HSPs. Cell Stress Chaperones 20, 605–620 (2015).

5. D. B. Mahat, H. H. Salamanca, F. M. Duarte, C. G. Danko, J. T. Lis, Mammalian Heat Shock Response and Mechanisms Underlying Its Genome-wide Transcriptional Regulation. Mol Cell 62, 63–78 (2016).

6. A. Currin et al., Highly multiplexed, fast and accurate nanopore sequencing for verification of synthetic DNA constructs and sequence libraries. Synth Biol (Oxf) 4, ysz025 (2019).

7. N. Pappireddi, L. Martin, M. Wühr, A Review on Quantitative Multiplexed Proteomics. Chembiochem 20, 1210–1224 (2019).

8. E. Reed et al., Assessment of a Highly Multiplexed RNA Sequencing Platform and Comparison to Existing High-Throughput Gene Expression Profiling Techniques. Front Genet 10, 150 (2019).

9. A. Keller, X. Tang, J. E. Bruce, Integrated Analysis of Cross-Links and Dead-End Peptides for Enhanced Interpretation of Quantitative XL-MS. Journal of proteome research 22, 2900–2908 (2023).

10. M. Barth, J. Bender, T. Kundlacz, C. Schmidt, Evaluation of NHS-Acetate and DEPC labelling for determination of solvent accessible amino acid residues in protein complexes. J Proteomics 222, 103793 (2020).

11. P. Limpikirati, T. Liu, R. W. Vachet, Covalent labeling-mass spectrometry with non-specific reagents for studying protein structure and interactions. Methods 144, 79–93 (2018).

12. S. Soni, P. Anand, Y. S. Padwad, MAPKAPK2: the master regulator of RNA-binding proteins modulates transcript stability and tumor progression. Journal of Experimental & Clinical Cancer Research 38, 1–18 (2019).

13. B. Goswami, S. Nag, P. S. Ray, Fates and functions of RNA-binding proteins under stress. WIREs RNA n/a, e1825.

14. S. Bresson et al., Stress-Induced Translation Inhibition through Rapid Displacement of Scanning Initiation Factors. Molecular Cell 80, 470–484.e478 (2020).

15. D. S. M. Ottoz, L. E. Berchowitz, The role of disorder in RNA binding affinity and specificity. Open biology 10, 200328 (2020).

16. S. Helder, A. J. Blythe, C. S. Bond, J. P. Mackay, Determinants of affinity and specificity in RNA-binding proteins. Current Opinion in Structural Biology 38, 83–91 (2016).

17. D. E. Draper, Themes in RNA-protein recognition. Journal of molecular biology 293, 255–270 (1999).

18. W. Miao, L. Li, Y. Wang, Identification of Helicase Proteins as Clients for HSP90. Analytical Chemistry 90, 11751–11755 (2018).

19. J. K. Flores, S. F. Ataide, Structural Changes of RNA in Complex with Proteins in the SRP. Frontiers in molecular biosciences 5, 7 (2018).

20. C. B. F. Andersen et al., Structure of the Exon Junction Core Complex with a Trapped DEAD-Box ATPase Bound to RNA. Science 313, 1968–1972 (2006).

21. G. Buchwald, S. Schüssler, C. Basquin, H. Le Hir, E. Conti, Crystal structure of the human eIF4AIII–CWC22 complex shows how a DEAD-box protein is inhibited by a MIF4G domain. Proceedings of the National Academy of Sciences 110, E4611–E4618 (2013).

22. W. Kabsch, C. Sander, Dictionary of protein secondary structure: Pattern recognition of hydrogen-bonded and geometrical features. Biopolymers 22, 2577–2637 (1983).

23. L. Ortega et al., 17-AAG improves cognitive process and increases heat shock protein response in a model lesion with Aβ25-35. Neuropeptides 48, 221–232 (2014).

24. T. Kijima et al., HSP90 inhibitors disrupt a transient HSP90-HSF1 interaction and identify a noncanonical model of HSP90-mediated HSF1 regulation. Scientific Reports 8, 6976 (2018).

25. H. H. Wippel, J. D. Chavez, A. D. Keller, J. E. Bruce, Multiplexed Isobaric Quantitative Cross-Linking Reveals Drug-Induced Interactome Changes in Breast Cancer Cells. Analytical Chemistry 94, 2713–2722 (2022).

26. K. E. Bohnsack, N. Kanwal, M. T. Bohnsack, Prp43/DHX15 exemplify RNA helicase multifunctionality in the gene expression network. Nucleic Acids Research 50, 9012–9022 (2022).

27. F. Hamann, M. Enders, R. Ficner, Structural basis for RNA translocation by DEAH-box ATPases. Nucleic Acids Res 47, 4349–4362 (2019).

28. K. Murakami, K. Nakano, T. Shimizu, U. Ohto, The crystal structure of human DEAH-box RNA helicase 15 reveals a domain organization of the mammalian DEAH/RHA family. Acta crystallographica. Section F, Structural biology communications 73, 347–355 (2017).

29. H. Walbott et al., Prp43p contains a processive helicase structural architecture with a specific regulatory domain. The EMBO journal 29, 2194–2204 (2010).

30. R. A. Becker, J. S. Hub, Continuous millisecond conformational cycle of a DEAH box helicase reveals control of domain motions by atomic-scale transitions. Communications Biology 6, 379 (2023).

31. R. Conde, Z. R. Belak, M. Nair, R. F. O’Carroll, N. Ovsenek, Modulation of Hsf1 activity by novobiocin and geldanamycin. Biochemistry and Cell Biology 87, 845–851 (2009).

32. G. E. Karagöz, S. G. D. Rüdiger, Hsp90 interaction with clients. Trends in Biochemical Sciences 40, 117–125 (2015).

33. F. H. Schopf, M. M. Biebl, J. Buchner, The HSP90 chaperone machinery. Nature Reviews Molecular Cell Biology 18, 345–360 (2017).

34. R. Y.-R. Wang et al., Structure of Hsp90–Hsp70–Hop–GR reveals the Hsp90 client-loading mechanism. Nature 601, 460–464 (2022).

35. C. M. Noddings, R. Y.-R. Wang, J. L. Johnson, D. A. Agard, Structure of Hsp90–p23–GR reveals the Hsp90 client-remodelling mechanism. Nature 601, 465–469 (2022).

36. R. C. Findly, R. J. Gillies, R. G. Shulman, In Vivo Phosphorus-31 Nuclear Magnetic Resonance Reveals Lowered ATP During Heat Shock of Tetrahymena. Science 219, 1223–1225 (1983).

## References

1. D. B. Mahat, H. H. Salamanca, F. M. Duarte, C. G. Danko, J. T. Lis, Mammalian Heat Shock Response and Mechanisms Underlying Its Genome-wide Transcriptional Regulation. Mol Cell 62, 63–78 (2016).

2. L. K. Pino, S. C. Just, M. J. MacCoss, B. C. Searle, Acquiring and Analyzing Data Independent Acquisition Proteomics Experiments without Spectrum Libraries. Molecular & Cellular Proteomics 19, 1088–1103 (2020).

3. B. C. Searle et al., Chromatogram libraries improve peptide detection and quantification by data independent acquisition mass spectrometry. Nature Communications 9, 5128 (2018).

4. S. I. Kovalchuk, O. N. Jensen, A. Rogowska-Wrzesinska, FlashPack: Fast and Simple Preparation of Ultrahigh-performance Capillary Columns for LC-MS. Mol Cell Proteomics 18, 383–390 (2019).

5. J. P. Mohr, P. Perumalla, J. D. Chavez, J. K. Eng, J. E. Bruce, Mango: A General Tool for Collision Induced Dissociation-Cleavable Cross-Linked Peptide Identification. Anal Chem 90, 6028–6034 (2018).

6. J. K. Eng, T. A. Jahan, M. R. Hoopmann, Comet: an open-source MS/MS sequence database search tool. Proteomics 13, 22–24 (2013).

7. J. D. Chavez, C. R. Weisbrod, C. Zheng, J. K. Eng, J. E. Bruce, Protein interactions, post-translational modifications and topologies in human cells. Mol Cell Proteomics 12, 1451–1467 (2013).

8. A. Keller, J. D. Chavez, J. E. Bruce, Increased sensitivity with automated validation of XL-MS cleavable peptide crosslinks. Bioinformatics 35, 895–897 (2019).

9. J. D. Chavez, A. Keller, J. P. Mohr, J. E. Bruce, Isobaric Quantitative Protein Interaction Reporter Technology for Comparative Interactome Studies. Anal Chem 92, 14094–14102 (2020).

10. J. Matthew Allen Bullock, J. Schwab, K. Thalassinos, M. Topf, The Importance of Non-accessible Crosslinks and Solvent Accessible Surface Distance in Modeling Proteins with Restraints From Crosslinking Mass Spectrometry. Molecular & cellular proteomics : MCP 15, 2491–2500 (2016).

11. W. Kabsch, C. Sander, Dictionary of protein secondary structure: Pattern recognition of hydrogen-bonded and geometrical features. Biopolymers 22, 2577–2637 (1983).

12. N. Yalamanchili et al., Distinct Cell Stress Responses Induced by ATP Restriction in Quiescent Human Fibroblasts. Frontiers in genetics 7, 171 (2016).

13. G. Grigolon et al., Grainyhead 1 acts as a drug-inducible conserved transcriptional regulator linked to insulin signaling and lifespan. Nat Commun 13, 107 (2022).

14. S. E. Nystrom, et al., JAK inhibitor blocks COVID-19 cytokine-induced JAK/STAT/APOL1 signaling in glomerular cells and podocytopathy in human kidney organoids. JCI insight 7, (2022).

15. S. Morikawa, L. Blacher, C. Onwumere, F. Urano, Loss of Function of WFS1 Causes ER Stress-Mediated Inflammation in Pancreatic Beta-Cells. Frontiers in endocrinology 13, 849204 (2022).

16. L. Ortega et al., 17-AAG improves cognitive process and increases heat shock protein response in a model lesion with Aβ25-35. Neuropeptides 48, 221–232 (2014).

17. T. Kijima et al., HSP90 inhibitors disrupt a transient HSP90-HSF1 interaction and identify a noncanonical model of HSP90-mediated HSF1 regulation. Scientific Reports 8, 6976 (2018).

18. H. H. Wippel, J. D. Chavez, A. D. Keller, J. E. Bruce, Multiplexed Isobaric Quantitative Cross-Linking Reveals Drug-Induced Interactome Changes in Breast Cancer Cells. Analytical Chemistry 94, 2713–2722 (2022).

19. W. Miao, L. Li, Y. Wang, Identification of Helicase Proteins as Clients for HSP90. Analytical Chemistry 90, 11751–11755 (2018).

